# Molecular hallmarks of heterochronic parabiosis at single cell resolution

**DOI:** 10.1101/2020.11.06.367078

**Authors:** Róbert Pálovics, Andreas Keller, Nicholas Schaum, Weilun Tan, Tobias Fehlmann, Michael Borja, James Webber, Aaron McGeever, Liana Bonanno, The Tabula Muris Consortium, Angela Oliveira Pisco, Jim Karkanias, Norma F. Neff, Spyros Darmanis, Stephen R. Quake, Tony Wyss-Coray

**Author notes:** These authors contributed equally (Róbert Pálovics, Andreas Keller, Nicholas Schaum, Weilun Tan). A full list of authors and affiliations appears in the online version of the paper.

## Abstract

Slowing or reversing biological ageing would have major implications for mitigating disease risk and maintaining vitality. While an increasing number of interventions show promise for rejuvenation, the effectiveness on disparate cell types across the body and the molecular pathways susceptible to rejuvenation remain largely unexplored. We performed single-cell RNA-sequencing on 13 organs to reveal cell type specific responses to young or aged blood in heterochronic parabiosis. Adipose mesenchymal stromal cells, hematopoietic stem cells, hepatocytes, and endothelial cells from multiple tissues appear especially responsive. On the pathway level, young blood invokes novel gene sets in addition to reversing established ageing patterns, with the global rescue of genes encoding electron transport chain subunits pinpointing a prominent role of mitochondrial function in parabiosis-mediated rejuvenation. Intriguingly, we observed an almost universal loss of gene expression with age that is largely mimicked by parabiosis: aged blood reduces global gene expression, and young blood restores it. Altogether, these data lay the groundwork for a systemic understanding of the interplay between blood-borne factors and cellular integrity.

---

Most ageing studies have focused on one or a few organs or cell types, with little temporal resolution. This has greatly limited our ability to interpret how and when ageing impacts interconnected organ systems. Recently, we performed a systematic characterization of ageing using bulk RNA-sequencing (RNA-seq) and single-cell RNA-sequencing (scRNA-seq) on dozens of mouse organs and cell types across the lifespan of the organism. We discovered both global and tissue/cell type-specific ageing signatures throughout the body^1–3^. But it remains unknown how, or if, rejuvenation paradigms affect these global ageing pathways in different cell types, or if nascent biochemical programs are instigated. The rational design of new therapeutics is therefore challenging.

One method of rejuvenation which has induced beneficial effects across organ systems is heterochronic parabiosis, in which a young and aged mouse share a common circulation. Phenotypes like cognition, muscle strength, and bone repair have all shown improvement through exposure to young blood in multiple laboratories^4^. And recently, epigenetic clock measurements in aged rats treated with young plasma demonstrated more youthful DNA-methylation profiles in multiple organs^5^. Parabiosis research has largely focused on age-related abundance changes to circulating proteins, and several proteins have been determined to mediate at least some of the observed effects^6–10^. However, such individual factors have yet to achieve robust rejuvenation throughout the body, likely in part due to an incomplete understanding of the effects of parabiosis on distinct organs and cells.

Here we attempt to address this question by performing Smart-seq2-based scRNA-seq of C57BL6/JN mice following 5 weeks of heterochronic parabiosis, when mice had reached 4 and 19 months of age (equivalent to humans aged around 25 and 65 years). Cells were captured via flow cytometry into microtiter plates from 13 organs: bladder, brain, brown adipose tissue (BAT, interscapular depot), diaphragm, gonadal adipose tissue (GAT, inguinal depot), limb muscle, liver, marrow, mesenteric adipose tissue (MAT), skin, spleen, subcutaneous adipose tissue (SCAT, posterior depot), and thymus (Fig. 1a,b, Extended Data Fig. 1a-d, Extended Data Tab. 1,2, n=1-4 individual mice per experimental group per organ). By integrating single-cell ageing data from the simultaneously collected *Tabula Muris Senis*, we were able to match cell type annotations per tissue based on k-nearest neighbors, and then compare parabiosis-mediated rejuvenation (REJ) and accelerated ageing (ACC) to normal ageing (AGE). Raw and annotated data are available from AWS (https://registry.opendata.aws/tabula-muris-senis/) and GEO (GSE132042).

## Cell type-specific differential gene expression

A fundamental unanswered question concerning parabiosis is which cell types are susceptible to accelerated ageing or rejuvenation (Fig. 1a). Out of a total of 13 tissues and >45,000 cells we were able to analyze differential gene expression in 54 cell types for ACC (isochronic young vs. heterochronic young) and 57 cell types for REJ (isochronic aged vs. heterochronic aged). Unexpectedly, we observe widespread transcriptomic changes, with 85 of 111 comparisons yielding >100 differentially expressed genes (DEGs) (Fig. 1c, Extended Data Fig. 2a-c), suggesting that nearly all cells are influenced by age-related changes in blood composition. The number of DEGs does not appear due to differences in cell number (Extended Data Fig. 2d-h) and differences between groups in percent mitochondrial genes, ribosomal genes, and ERCCs are not evident (Extended Data Fig. 1e-g). Furthermore, permuting the experimental groups within each cell type resulted in fewer than 100 DEGs in 104 cases out of 111, indicating that the hundreds to thousands of DEGs resulting from heterochronic parabiosis are not random and likely the result of biology (Extended Data Fig. 3).

**Fig. 1.**
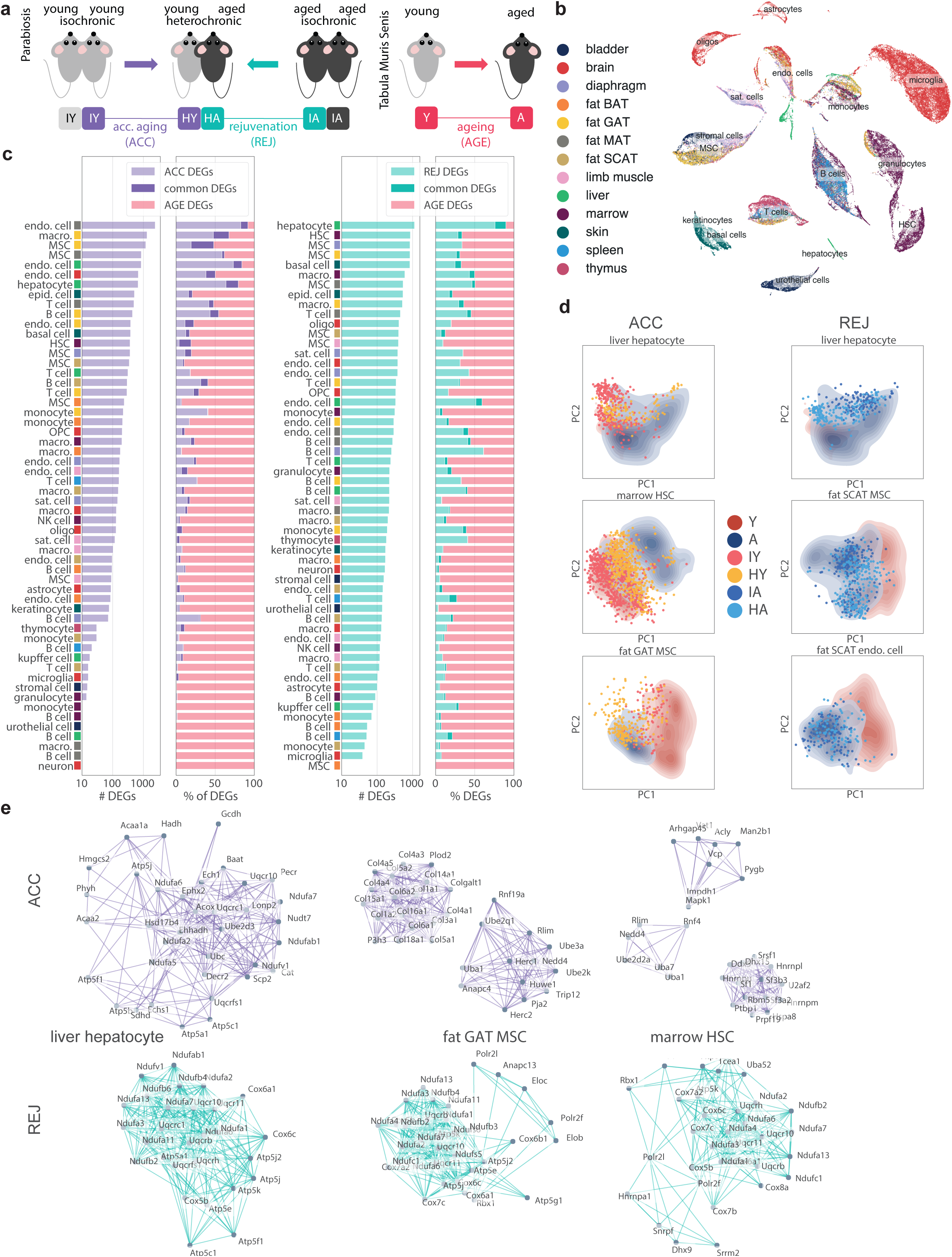
Cell type-specific differential gene expression. **a**, Experimental outline. FACS/Smart-seq2-based scRNA-seq data was collected from male isochronic and heterochronic pairs (n=1-2 individual mice per group; 3-months and 18-months-old), and integrated with FACS/Smart-seq2-based scRNA-seq data from *Tabula Muris Senis* male mice (n=4 3-month-old mice; n=6 18-24-month-old mice). IY: isochronic young. HY: heterochronic young. HA: heterochronic aged. IA: isochronic aged. Y: young. A: aged. ACC: accelerated ageing. REJ: rejuvenation. AGE: ageing. **b**, Uniform manifold approximation and projection (UMAP) based on the first 16 principle components of all parabiosis cells (n=45,331 cells from 13 tissue types). **c**, Cell types ranked by the number of differentially expressed genes (DEGs) for ACC (left) and REJ (right). The percentage of DEGs overlapping with those or normal ageing (AGE) is indicated. The number of cells used for differential expression is in Extended Data Fig. 2. Differential gene expression was conducted on the CPM normalized and log-transformed count matrix (p<0.01, eff>0.6, |log_2_FC|>0.5). **d**, The first two principal components (PC1; PC2) for select parabiosis cell types, with the corresponding cells from *Tabula Muris Senis* as background heatmaps. PCA was conducted on DEGs as in (**c**), after pre-selecting the strongest ageing genes (p<0.01, eff>0.6, |log_2_FC |>0.5). **e**, Densest STRING subnetwork of the top DEGs that are consistent with AGE DEGs for select cell types. ACC (top), REJ (bottom). STRING links with >0.9 confidence (scale from 0-1) are queried, and the densest k-core subgraph is shown.

**Fig. 2.**
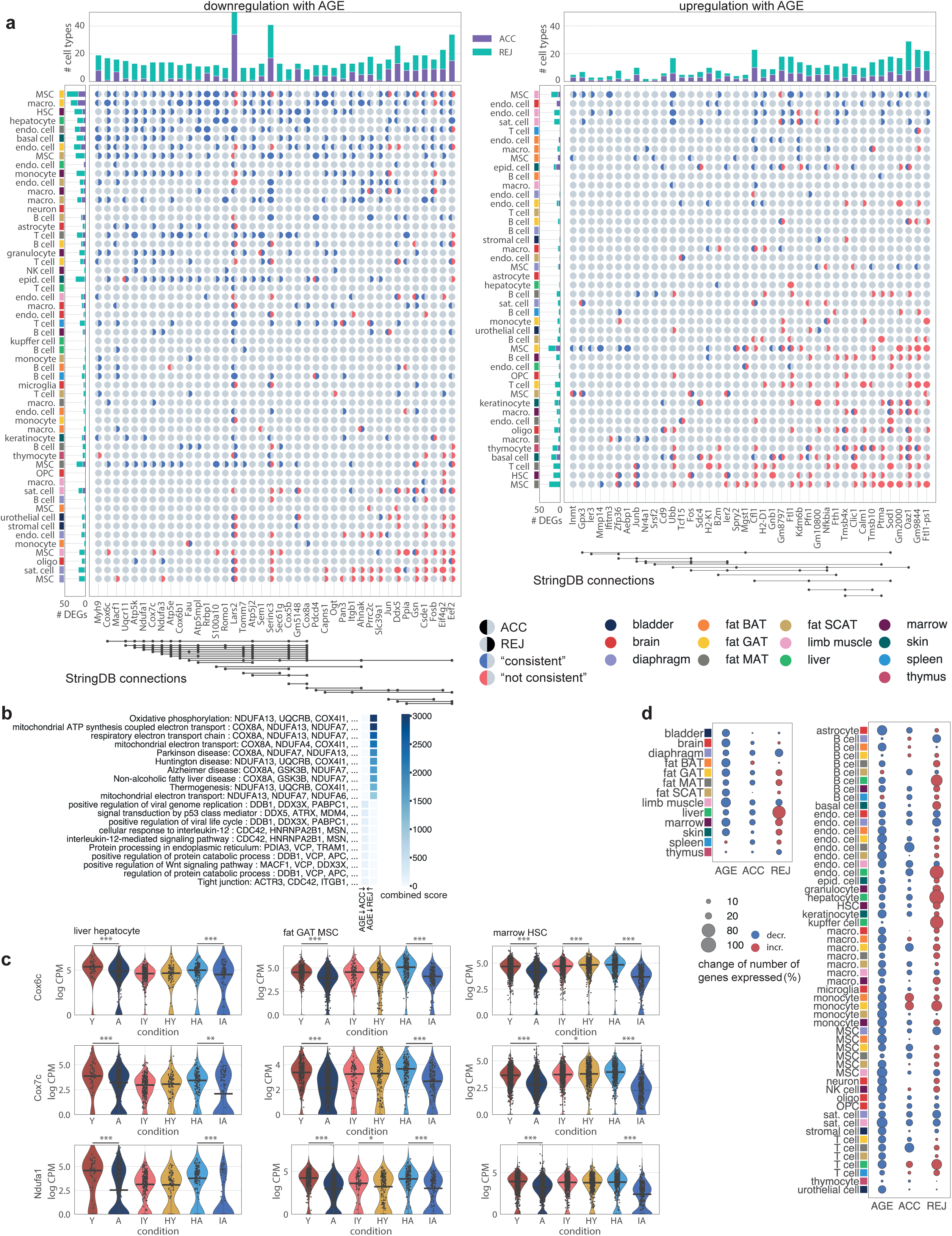
Young blood reverses mitochondrial & global gene expression loss. **a**, DEGs downregulated (left) or upregulated (right) with age that are most frequently rescued (i.e. “consistent”) across cell types by parabiosis. “Not consistent” indicates if the direction of change is identical for both ageing and parabiosis. Columns and rows are sorted by cases of “consistent” minus “not consistent”. Top: the number of cell types for rejuvenation (REJ) and accelerated ageing (ACC) for which each gene is differentially expressed (“consistent” + “not consistent”). Left: the number of total DEGs per cell type (“consistent” + “not consistent”). Bottom: STRING connections between top genes. STRING links with >0.9 confidence (scale from 0-1) are queried, and the densest k-core subgraph is shown. **b**, Most enriched pathways (GO Biological Process and KEGG) among the top 200 ACC/REJ genes consistently changing with ageing downregulation. **c**, Violin plots for liver hepatocytes, GAT MSCs and marrow HSCs of select genes encoding proteins of the electron transport chain. **d**, Relative change of the mean number of genes expressed for each cell type (left) and combined cell types for each tissue (right).

**Fig. 3.**
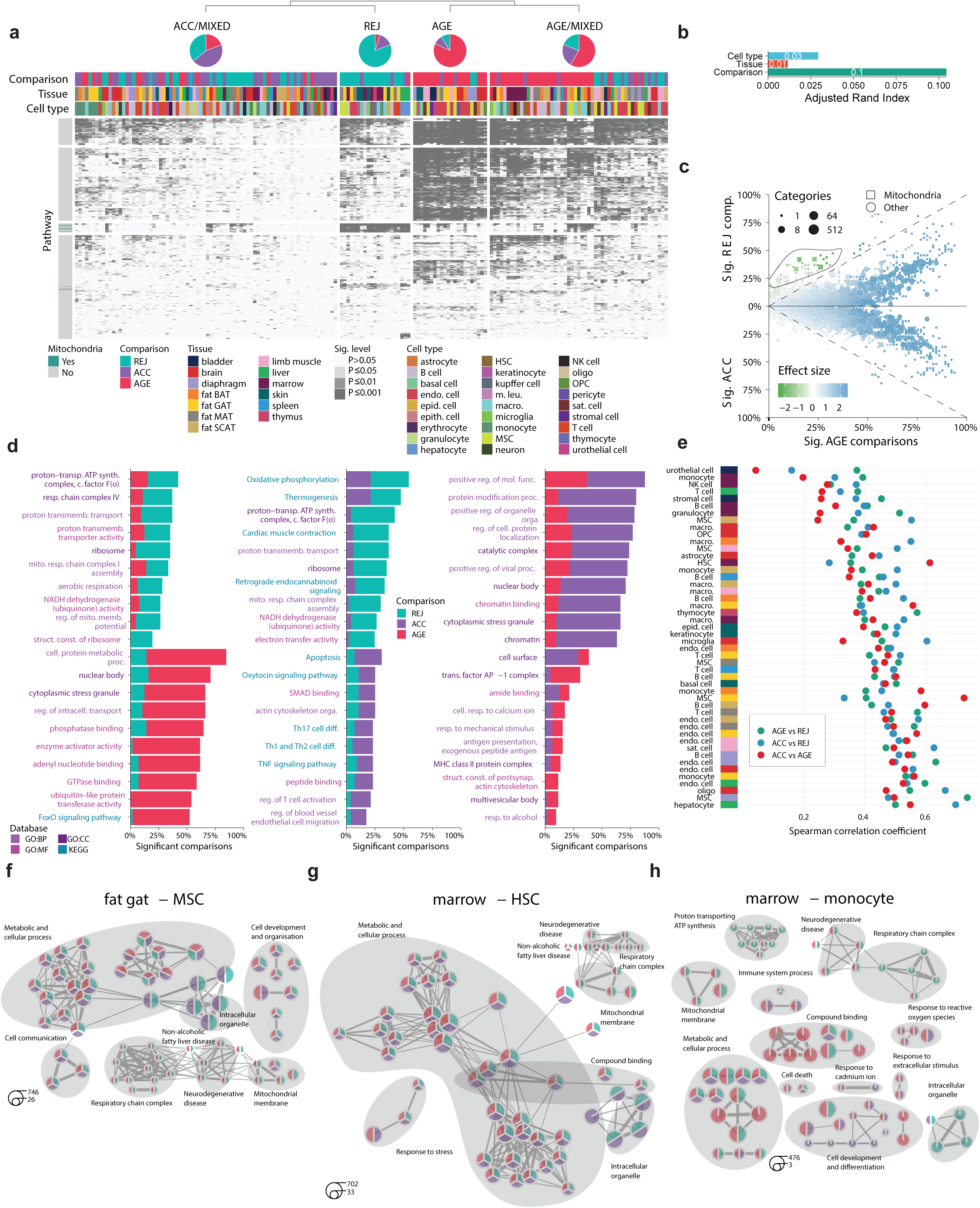
Structured responses to parabiosis. **a**, Pathway enrichment and clustering for all cell types for ageing, accelerated aging, and rejuvenation (GO and KEGG). Four modules are evident, each described by its proportion of each of the three comparisons (top). Mitochondrial pathways highlighted in teal. **b**, The adjusted rand index (ARI) for the four clusters. **c**, The percentage and effect size of significant tissues and cell types per pathway. **d**, For each pairwise comparison between ageing, rejuvenation, and accelerated ageing, the top pathways are indicated, ranked by the percentage of cell types in which they emerge. **e**, Spearman correlation coefficient of DEGs within each cell type between comparisons (ageing, rejuvenation, accelerated ageing). Each block indicates a cell type within the designated tissue. Top pathways for GAT MSCs (**f**), marrow HSCs (**g**) and marrow monocytes (**h**). The proportion of each pathway derived from each comparison is indicated via pie chart, and related pathways are grouped into modules.

Most prominently, hepatocytes exposed to young blood show 1,000 DEGs, with heterochronic aged hepatocytes undergoing a clear shift toward young in principal component analysis (Fig. 1c, d). In fact, young hepatocytes exposed to aged blood undergo marked ageing, with more than 600 DEGs. Considering the liver is the most highly perfused organ and the major source of plasma proteins, these cells appear to be exceptionally responsive to age-related changes in the systemic environment. Befittingly, these were one of the first cell types described to undergo rejuvenation from exposure to young blood^11^.

Hepatocytes are perhaps only surpassed in their proximity to blood by those cells that line blood vessels themselves – endothelial cells (ECs). With 2,429 DEGs after exposure to aged blood, ECs of the viscerally located MAT represent the most substantial transcriptomic response among all cells. ECs from the brain, liver, and visceral GAT all feature among the top 11 accelerated ageing cell types, with 300–1,000 DEGs, suggesting that continuous and direct exposure to the aged circulatory system induces strong transcriptomic changes. With 80-2,432 DEGs each due to young or aged blood, ECs across all tissues seem susceptible to blood-borne influences, albeit with tissue-specificity, perhaps due to differences in perfusion, differences in cell intrinsic programs, or influence from parenchymal cells. Recently, transfused aged plasma was shown to recapitulate transcriptomic ageing of hippocampal ECs, and young plasma reversed aspects of ageing, especially in capillary ECs^12^.

Just like ageing of blood vessels, ageing of fat tissues substantially contributes to disease risk and declining health. Specifically, visceral adipose tissues undergo some of the earliest and most dramatic transcriptomic changes with age^2^, and the expansion and inflammation of visceral fat is especially detrimental. In addition to strong parabiosis-mediated changes in MAT and GAT endothelial cells, mesenchymal stromal cells (MSCs) in both tissues display large numbers of DEGs, and principle component analysis reveals marked shifts after exposure to differentially aged blood (Fig. 1d). In fact, MSCs from adipose tissues exhibit hundreds of DEGs in both young and aged heterochronic parabionts. In line with recent findings that the pro-ageing systemic protein CCL11^6^ is produced in visceral adipose tissue^13^, Ccl11 and other age-related genes encoding plasma proteins like Chrdl1 and Hp are within the first two principal components for GAT MSCs (Extended Data Fig. 4a-e), indicating that these cells may be contributors to ageing of the systemic environment. As well, preadipocytes within the MSC population share many characteristics with tissue-resident macrophages, and GAT macrophages actually feature among the top cell types changed with parabiosis (Fig. 1c).

**Fig. 4.**
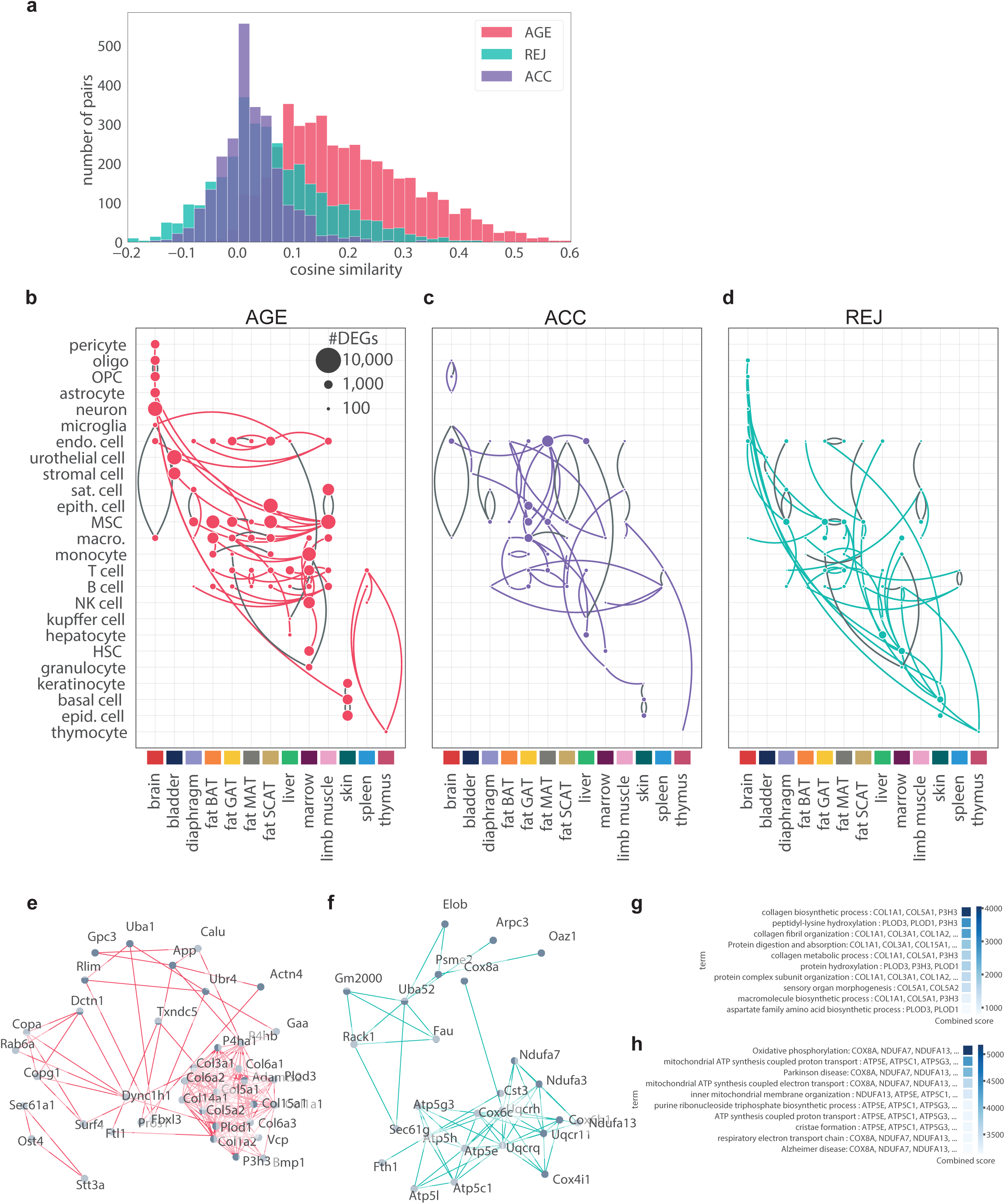
Coordinated, organism-wide cellular responses to ageing and parabiosis. **a**, Histogram of cosine similarity of ageing, accelerated ageing, and rejuvenation gene signatures between each pairwise comparisons of cell types. **b-d**, Based on the cosine similarities from (**a**), each cell type is connected to its most similar cell type. Grey indicates non-unique connections. **e**, STRING network of DEGs common to MSCs from GAT, MAT, SCAT, BAT, limb muscle, and diaphragm. **f**, STRING network of DEGs common to MSCs (GAT and MAT), hepatocytes, basal and epidermal cells (skin), and HSCs and macrophages (marrow). STRING links with >0.9 confidence (scale from 0-1) are queried, and the densest k-core subgraph is shown. **g-h** Most enriched pathways (GO Biological Process and KEGG) among the nodes of the networks shown in (**e-f**).

Immune cell accumulation in adipose depots is a fundamental feature of ageing, and indeed most types, including T cells, B cells, neutrophils, and plasma cells, accrue across diverse organs^2,14^. It is interesting that tissue-resident immune cells of both the lymphoid lineage (T, B, NK cells) and myeloid lineage (monocytes & macrophages) seem liable to the effects of parabiosis, as do their marrow-resident precursors, hematopoietic stem cells (HSCs; Fig. 1b). In fact, 1,000 HSC genes are altered by young blood, perhaps indicating a tight-knit relationship between ageing of the immune system and changes in blood composition. Previously, heterochronic transplantation of marrow or HSCs in mice has been shown to affect (modulate) a variety of phenotypes^15–18^. Most recently, aged HSCs were found to induce circulating cyclophilin A, encoded by Ppia^19^, a gene ranked among the top differentially expressed across cell types exposed to aged blood (Extended Data Fig. 5). Yet here, heterochronic aged HSCs do not appear to shift via PCA, suggesting that young blood acts primarily on non-ageing pathways.

We therefore asked if parabiosis induces reversal or acceleration of ageing pathways, or if novel genes are invoked. After integrating FACS-Smart-seq2 data from >37,000 *Tabula Muris Senis* cells, we found that for a number of cell types, most notably endothelial cells and MSCs, the effects of parabiosis are equal to - or even much more pronounced than - ageing, suggesting that these cells are particularly susceptible to changes in plasma composition with age. In three cases, a substantial number of accelerated ageing DEGs overlap with those of normal ageing: 60% for GAT MSCs, 80% for HSCs, and 84% for oligodendrocytes (Fig. 1c). Endothelial cells from a variety of tissues are also consistently among the top cell types with the most parabiosis DEGs in common with ageing. This suggests that a significant part of ageing of these cells may be attributed to ageing of the systemic environment. Overall, aged blood induces changes more akin to ageing than young blood, as can be seen by the larger proportion of overlapping DEGs for many cell types (Fig. 1c). However, rejuvenation appears to be a much more concerted process: the core network of ageing DEGs rescued by rejuvenation consists of mitochondrial electron transport chain genes for multiple cell types (Fig. 1e).

As well, there are numerous instances where accelerated ageing and rejuvenation have little to no overlap with ageing DEGs. The reason for these discordant results is currently unknown, but it could be that ageing of these cells is influenced more by other factors, masking subtler effects caused by an altered systemic circulation. Overall, these data indicate that nearly all cell types are amenable to reformation via changes to blood composition, even those not directly exposed to blood. Furthermore, it appears that ageing of certain cell types - especially HSCs which give rise to circulating and tissue-resident immune cells - is heavily influenced by the systemic milieu.

## Young blood reverses mitochondrial & global gene expression loss

While ageing is in part manifested differently across tissues and cell types, the substantial overlap in ageing signatures suggests that targeting common molecular pathways – by modifying blood composition, for example – could slow or reverse harmful changes throughout the body. We therefore aimed to determine if parabiosis reverses or accelerates ageing gene expression signatures that are common to multiple cell types and tissues. We first selected genes differentially expressed in the most cell types for both parabiosis and ageing, and indicated agreement with ageing based on the direction of change (Fig. 2a). Foremost is the pronounced disparity between genes upregulated and those downregulated during ageing. While upregulated genes appear largely sporadic, downregulated genes show considerable agreement with parabiosis and enrichment for biological pathways. Furthermore, permuting the experimental groups within each cell type resulted in almost no overlap with ageing DEGs (Extended Data Fig. 3). Most conspicuously, across a range of cell types and tissues, exposure to young blood increases the expression of genes encoding electron transport chain subunits like *Cox6c, Cox7c, Ndufa1, Ndufa3, Atp5k*, and *Uqcr11*, reversing the loss of expression in normal ageing (Fig. 2a,c). In fact, oxidative phosphorylation and the electron transport chain are the top enriched pathways (Fig. 2b), and of the parabiosis DEGs that agree most consistently with ageing.

The loss and dysregulation of mitochondrial function is one of the most ubiquitous and drastic mammalian ageing hallmarks^2,10^, so the widespread rejuvenation observed here hints that systemic restoration may be possible through manipulation of the systemic environment. Indeed, rejuvenation of such genes is visible for a variety of cell types, from HSCs and hepatocytes to endothelial cells and immune cells across tissues (Fig. 2a). Notably, such signatures are absent in certain cell types (Extended Data Fig. 6), including brain endothelial cells (BECs), which have been observed to undergo increased expression of electron transport chain genes with age^12^. This effect is replicated by exposure to aged mouse plasma, and reversed by exposure to young mouse plasma in vivo^12^. Such an exception to the global pattern could prove useful for elucidating the mechanism of crosstalk between blood factors and mitochondrial function.

Restoration of mitochondrial gene expression is but one part of a more global response to young blood: not only is gene expression loss with age evident in nearly every cell type, but this is mimicked by accelerated ageing and reversed by rejuvenation (Fig. 2d). This supports a fundamental role for transcriptional regulation itself in ageing and rejuvenation paradigms.

## Structured responses to parabiosis

In spite of the striking rescue to age-related gene expression loss – and specifically to genes encoding proteins of the mitochondrial electron transport chain - ageing-independent pathways may also contribute to the parabiosis-mediated functional improvements observed throughout the body. To investigate the molecular hallmarks induced by parabiosis in each cell type, we performed unbiased pathway analysis for each cell type in response to ageing and a young or aged circulatory environment (AGE, REJ, ACC). We identified 4 major pathway clusters (Fig. 3a) which were largely driven by the environment the cells are exposed to rather than inherent cell or tissue type, as indicated by the adjusted rand index score (Fig. 3b). This clustering highlights the widespread and comparatively strong influence of ageing, but it also reveals that ACC largely affects the body either through systemic changes that mimic those of ageing, or sporadic, cell type-specific effects. On the other hand, cluster 2 indicates ageing- and ACC-independent pathways, suggesting rejuvenation invokes novel biology, whereas the remaining REJ pathways overlap with ageing or ACC. These data are confirmed when comparing the percentage of cell types enriched for specific pathways (Fig. 3c), with ageing dominating pathway analysis, and REJ effects that either act outside ageing pathways or act to oppose ageing pathways. Foremost, pathway analysis reveals that the electron transport chain is widely altered across ageing and REJ, suggesting enhanced metabolic activity in heterochronic aged parabionts (Fig. 3c,d).

We next sought to determine in which cell types parabiosis-mediated REJ or ACC pathways most closely agree or disagree with ageing. For each cell type we calculated the spearman correlation coefficient between pathways for ageing & REJ, ageing & ACC, and REJ & ACC (Fig. 3e). With a correlation of 0.73, GAT MSCs display a highly similar transcriptional response to ageing and aged blood, as do HSCs (ρ=0.61). The absence of mitochondrial electron transport genes common to these two groups is notable. Such genes commonly arise and overlap between ageing and rejuvenation (Fig. 3f,g), even in cell types for with ACC correlates more strongly with ageing than REJ does. This suggests that young blood is a potent instigator of mitochondrial function, while aged blood itself contributes little to the age-related decline. In fact, mitochondrial genes arise even for cell types in which age-related decline is not evident, like marrow monocytes (Fig. 3h), supporting the notion that young blood may indeed broadly enhance mitochondrial function.

There are also cell types for which rejuvenation is highly correlated with ageing, exemplified by MSCs from the diaphragm (ρ=0.74). In fact the same cell type, present in different organs, often shows highly divergent responses to ageing, accelerated ageing, and rejuvenation, indicating that the immediate environment in which a cell resides often exerts more influence than circulating factors. The consistent exception is endothelial cells, which show high correlation between AGE, ACC, and REJ, regardless of tissue of origin. We conclude that these cells are especially susceptible to influences from the systemic environment due to their continuous exposure to blood.

## Coordinated, organism-wide cellular responses to ageing and parabiosis

In order to appreciate the overarching effects of ageing and parabiosis organism-wide, we asked if individual cell types throughout the various tissues of the mouse show similar or discordant transcriptional responses to aging and parabiosis. For all pairwise comparisons between cell types, we computed the cosine similarity of their respective DEGs for ageing, rejuvenation, and accelerated ageing (Fig. 4a). While the highest similarities are evident for ageing, the transcriptomic signature of rejuvenation elicited by young blood also shows considerable conservation between cell types. Such commonalities are lacking for accelerated ageing, in agreement with the divergent pathways arising for top ACC DEGs (Fig. 3d). To determine which groups of cell types are responsible for the ageing similarity signature, we plotted the closest connection for each cell type (Fig. 4b). Remarkably, ageing instigates coordinated transcriptomic changes with high similarity within some tissues, most notably brain, skin, and marrow, yet clearly distinct signatures between tissues overall, suggesting that local pro-ageing factors or programs may govern ageing of these tissues. Equally exciting, we discovered that cell types, such as endothelial cells, MSCs, and immune cells share transcriptional programs of ageing across vastly different and distant tissues, possibly reflecting cell intrinsic transcriptional programs of ageing. Indeed, for mesenchymal stromal cells across four adipose tissues and two skeletal muscle types, the loss of collagen gene expression forms a core node (Fig. 4e,g). In the context of immune cells, it has been speculated that infiltration of these cells may lead to “spreading” of ageing in invaded tissues through secreted factors^2,20^. Future studies may explore the basis of cellular “hubs” which are transcriptionally related to many cell types - e.g. monocytes of marrow, ECs of SCAT - while other cell types are less connected.

A similar analysis of parabiosis shows that an aged circulation mimics, in part, the tissue and cell type specific transcriptional similarities, but they are overall less pronounced, and many seem to disappear (Fig. 4c). Intriguingly, while skin and marrow maintain solid tissue-wide cellular transcriptomes following exposure to young blood – albeit different from those observed with aging – many new transcriptional similarities emerge across cell types and tissues (Fig. 4d). Most notably, REJ triggers similar transcriptional signatures across highly divergent cell types. For example, the mitochondrial electron transport gene node emerges once again as a core rejuvenation network, and is especially strong between MSCs (GAT, MAT), hepatocytes, basal and epidermal cells from skin, and HSCs and macrophages from marrow (Fig. 4f,h).

## Discussion

Our dataset provides a first look into the transcriptomic effects of heterochronic parabiosis at single-cell resolution. Continuous exposure to differentially aged blood alters the transcriptomic landscape across cell types, and we discovered that particular cell types - namely MSCs, ECs, HSCs, and hepatocytes - are especially susceptible to gene expression changes. While the effects of aged blood tend to accelerate normal ageing changes, young blood both reverses age-related profiles and initiates novel pathways. Systemic rejuvenation of genes encoding components of the electron transport chain is especially striking, as is the reversal of global gene expression loss with age. Together, these findings reveal the molecular details of how ageing and parabiosis trigger highly complex global responses across the organism, some of which are tissue-specific and some cell type-specific, likely reflecting a sophisticated combination of cellular, local, and systemic transcriptional cues. These newly discovered transcriptional programs shared between cell types in response to the three chronogenic environments suggest possible avenues for therapeutic interventions. Finally, heterochronic parabiosis represents only one rejuvenation paradigm, and organism-wide analysis of other interventions, such as was recently conducted for caloric restriction in rats^14^, may help uncover complimentary treatments able to more comprehensively target ageing hallmarks throughout the body.

## Methods

### Experimental procedures

#### Parabiosis and organ collection

3-month-old and 18-month-old male C57BL/6JN mice were shipped from the National Institute on Ageing colony at Charles River (housed at 19–23□°C) to the Veterinary Medical Unit (VMU; housed at 20–24□°C)) at the VA Palo Alto (VA). At both locations, mice were housed on a 12 h/12 h light/dark cycle and provided with food and water ad libitum. The diet at Charles River was NIH-31, and at the VA VMU was Teklad 2918. Littermates were not recorded or tracked, and mice were housed at the VA VMU for no longer than 2 weeks before surgery.

Parabiosis via the peritoneal method was accomplished by suturing together the peritoneum of adjacent flanks, forming a continuous peritoneal cavity. To promote coordinated movement, adjacent knee joints and elbow joints were joined with nylon monofilament sutures. Skin was joined with surgical autoclips. All procedures were conducted with aseptic conditions on heating pads, with mice under continuous isoflurane anesthesia. To prevent infection, limit pain, and promote hydration, mice were injected with Baytril (5 ug/g), Buprenorphine, and 0.9% (w/v) sodium chloride, as described previously^4,21^. Pairs remained together for 5 weeks prior to organ collection.

After anaesthetization with 2.5% v/v Avertin at 8:00, mice were weighed, shaved, and blood was drawn via cardiac puncture before transcardial perfusion with 20 ml PBS. Mesenteric adipose tissue was then immediately collected to avoid exposure to the liver and pancreas perfusate, which negatively affects cell sorting. Isolating viable single cells from both the pancreas and the liver of the same mouse was not possible; therefore only one was collected from each mouse. Whole organs were then dissected in the following order: large intestine, spleen, thymus, trachea, tongue, brain, heart, lung, kidney, gonadal adipose tissue, bladder, diaphragm, limb muscle (tibialis anterior), skin (dorsal), subcutaneous adipose tissue (inguinal pad), brown adipose tissue (interscapular pad), aorta and bone marrow (spine and limb bones). Organ collection concluded by 10:00. After single-cell dissociation as described below, cell suspensions were used for FACS of individual cells into 384-well plates. All animal care and procedures were carried out in accordance with institutional guidelines approved by the VA Palo Alto Committee on Animal Research.

### Sample size, randomization and blinding

No sample size choice was performed before the study. Blinding was not performed: the authors were aware of all data and metadata-related variables during the entire course of the study.

### Tissue dissociation and sample preparation

All tissues were processed as previously described^3^.

## Single-cell methods

All protocols used in this study are described in detail elsewhere^1,3^. These include: preparation of lysis plates; FACS sorting; cDNA synthesis using the Smart-seq2 protocol^22,23^; library preparation using an in-house version of Tn5^24,25^; library pooling and quality control; and sequencing. For further details please refer to https://www.protocols.io/view/smartseq2-for-htp-generation-of-facs-sorted-single-2uwgexe.

## Computational methods

### Data extraction

We unified these data with scRNA-seq profiles of cells from young (3-month-old males) and aged (combined 18-month-old & 24-month-old males) mice from the *Tabula Muris Senis* Smart-seq2 data^2,3^. All subsequent data processing and analysis is conducted on this unified dataset.

## Quality control

We applied standard filtering rules following the guideline of Luecken et al.^26^. We discarded cells with (1) fewer than 500 genes or (2) less than total 5,000 reads or (3) more than 30% ERCC reads or (4) more than 10% mitochondrial reads or (5) more than 10% ribosomal reads. Counts were then CPM scaled and log-normalized for downstream analysis. Analysis was implemented with Gseapy 0.10.1, Matplotlib 3.3.2, Networkx^27^ 2.5, Numpy v1.18.1, Pandas v1.0.1, Scanpy^28^ v1.4.4, Scikit-learn^29^ v0.22.1, and Seaborn 0.11.0 packages.

### Cell type annotations

We grouped the data based on tissue of origin and computed 32 principal components (PCA) of the normalized data for each tissue. We embedded the cells in a 32-dimensional latent space using these PCA components and then identified their k=20 nearest neighbors from the *Tabula Muris Senis* data. We then applied majority voting to define the type each cell from the parabionts. In other words, we calculated the most frequent cell type among the cell’s neighbors from *Tabula Muris Senis* and used this to annotate the cell. Note that *Tabula Muris Senis* includes some highly specific cluster annotations and we joined some of these to achieve more robust results, e.g. we merged all the T cell subclusters. These merging rules can be found in Extended Data Table 3. Finally, to visualize the cell clusters we computed UMAP embeddings^30^. We ran the DBSCAN clustering algorithm (eta=0.8) on the UMAP embeddings in order to identify groups of cells that are not present in both datasets. We discarded clusters of cells from the analysis that were only present in TMS. To get a global picture of the final dataset we repeated the PCA and UMAP computations over all cells together. Analysis was implemented in Python 3.8.3 with Gseapy 0.10.1, Matplotlib 3.3.2, Networkx 2.5, Numpy v1.18.1, Pandas v1.0.1, Scanpy v1.4.4, Scikit-learn v0.22.1, and Seaborn 0.11.0 packages.

### Differential gene expression

We systematically analyzed parabiosis signatures across 3 comparisons (Y-O, IY-HY, IO-HO) within each identified cell type. We conducted single-cell differential gene expression for the 3 comparisons within each cell type separately. Specifically, we computed standard log2-fold changes as well as the non-parametric unpaired Wilcoxon–Mann–Whitney test^31^ for each gene. Finally, we identified genes differentially expressed with effect size>0.6, p-value<0.01 and |log_2_FC|>0.5. Note that the effect size of the Wilcoxon–Mann–Whitney test is the AUC metric, frequently used in case of large datasets since it is not sensitive to the sample size. Hence filtering for this metric is especially important as single-cell data often contains large sample sizes with thousands of cells per condition. We discarded genes used for QC filtering from the DGE analysis. Analysis was implemented in Python 3.8.3 with Gseapy 0.10.1, Matplotlib 3.3.2, Networkx 2.5, Numpy v1.18.1, Pandas v1.0.1, Scanpy v1.4.4, Scikit-learn v0.22.1, and Seaborn 0.11.0 packages.

### Pathway analysis

Over-representation analysis was performed using GeneTrail 3^32^ for all significantly deregulated genes in ageing, accelerated ageing and rejuvenation, per tissue and cell type using the categories of Gene Ontology^33^ and KEGG pathways^34^. P-values were adjusted for multiple testing per database using the Benjamini-Hochberg procedure^35^. Depleted categories were not considered. Results were analyzed with the programming language R 4.0.2. To generate the enrichment heatmap the 30 most enriched categories of each comparison were extracted. The columns of the enrichment matrix were clustered with hierarchical clustering using Ward’s clustering criterion and Euclidean distance, based on the discretized P-values (<0.05, <0.01, <0.001). The clustering was cut at a height of 4. The rows were clustered with complete linkage and Euclidean distance. The heatmap was plotted with the ComplexHeatmap^36^ (2.4.2) R package. To determine the major clustering factor among the comparison, tissues or cell types, we computed the adjusted rand index (ARI) using the aricode R package (1.0.0) and plotted them as bar plot with ggplot2^37^ (3.3.2). For determining the most different pathways per comparison, we filtered similar terms using the GOSemSim R package (2.14.0) according to the Jiang measure with a cutoff at a similarity of 0.7, and excluded KEGG disease pathways. We computed for every setup comparison the per tissue and cell type similarity of the determined enrichment P-values on the negative log10 transformed values by using Spearman’s correlation coefficient. Pathway and gene set networks were generated for each tissue and cell type using the 30 most significant enrichments and plotted with igraph^38^ (1.2.5), ggraph (2.0.3), and scatterpie (0.1.5).

### PCA analysis of responding cell types

For each cell type showing strong response to parabiosis, first we selected *ageing* genes that were differentially expressed with effect size>0.6, p-value<0.01 and |log_2_FC |>0.5 in case of the Y-O comparison. We then carried out principal component analysis (PCA) across these ageing genes. In our results we show the 1st and 2nd PCA components of each cell from the parabionts. We present the ageing signal in the background with kernel density estimation. Analysis was implemented in Python 3.8.3 with Gseapy 0.10.1, Matplotlib 3.3.2, Networkx 2.5, Numpy v1.18.1, Pandas v1.0.1, Scanpy v1.4.4, Scikit-learn v0.22.1, and Seaborn 0.11.0 packages.

### Ageing and rejuvenation similarity analysis

We base these analyses on the differential gene expression results. We define similarities for the 3 comparisons (Y-O, IY-HY, IO-HO) separately. First, we select genes that are differentially expressed with effect size>0.6, p-value<0.01 and |log_2_FC|>0.5. Next, we take the vectors indicating the direction of the expression changes across these genes in case of each cell type (+1 up, 0 no change, -1 down). We compute then the cosine similarities of those vectors to define pairwise similarities between the cell types. We present the structure of these similarity networks in our results. Analysis was implemented in Python 3.8.3 with Gseapy 0.10.1, Matplotlib 3.3.2, Networkx 2.5, Numpy v1.18.1, Pandas v1.0.1, Scanpy v1.4.4, Scikit-learn v0.22.1, and Seaborn 0.11.0 packages.

### STRING network analysis

For each set of DEGs of interest, first we queried the STRING database for links with >0.9 confidence. Next we selected the densest component of the network with no more than 40 DEGs within it. Selection was done by k-core decomposition: we recursively pruned the network to select its subnetwork where each node’s degree is at least k. We set k in order to find the densest core with no more than 40 DEGs. We used the k-core implementation of networkx 2.5 python package.

## Code Availability

All code used for analysis will be available upon publication.

### Data Availability

Raw and annotated data are available on AWS (https://registry.opendata.aws/tabula-muris-senis/) and GEO (GSE132042).

## Acknowledgements

We thank the members of the Wyss-Coray laboratory and the Chan-Zuckerberg Biohub for feedback and support. Funding for library preparation, sequencing, and AWS time was provided by the Chan Zuckerberg Biohub. Additional funding includes the Department of Veterans Affairs (BX004599 to T.W.-C.), the National Institute on Aging (R01-AG045034 and DP1-AG053015 to T.W.-C.), the NOMIS Foundation (T.W.-C.), The Glenn Foundation for Medical Research (T.W.-C.), and the Wu Tsai Neurosciences Institute (T.W.-C.). This work was supported by the National Institute of Aging and the National Institutes of Health under award number P30AG059307.

## Author Contributions

R.P., A.K., N.S., and W.T. contributed equally. N.S., S.R.Q., and T.W.-C. conceptualized the study. R.P., A.K., N.S., T.F., and T.W.-C. conceptualized the analysis. R.P., A.K., and T.F., conducted the analysis. N.S. and L.B. performed parabiosis surgeries. The *Tabula Muris Consortium* processed organs and captured cells. W.T. and M.B. conducted cDNA and library preparation. R.P. created the web browser. W.T. and M.B. performed sequencing and library quality control. W.T., M.B., A.O.P., J.W., and A.M. processed raw sequencing data. R.P., A.K., N.S., S.R.Q., and T.W.-C., wrote and edited the manuscript. T.W.-C., S.R.Q., S.D., N.F.N., J.K., and A.O.P supervised the work.

**Supplementary Information** is available in the online version of the paper.

## Author Information

Reprints and permissions information is available at www.nature.com/reprints. The authors declare no competing financial interests. Readers are welcome to comment on the online version of the paper. Correspondence to, and.

## Reviewer Information

*Nature* thanks the anonymous reviewers for their contributions to the peer review of this work.

**Extended Data Fig. 1.**
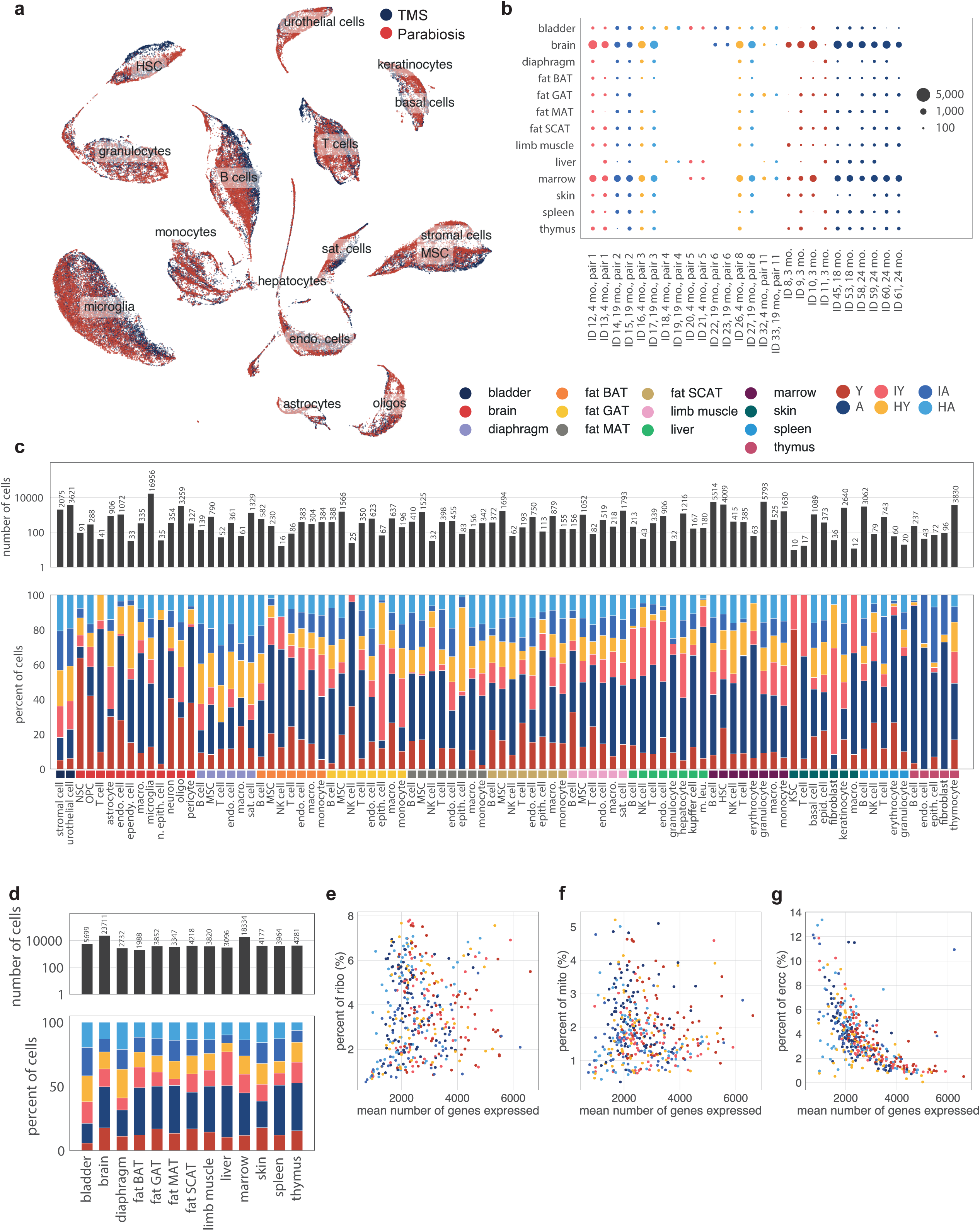
**a**, Uniform manifold approximation and projection (UMAP) based on the first 16 principle components of all parabiosis and TMS cells (n= 83,277 cells from 13 tissue types). **b**, Number of cells per tissue and mouse. **c**, Total number of cells per cell type (top), fraction of cells covering each of the 6 experimental conditions per cell type. **d**, Total number of cells per tissue (top), fraction of cells covering each of the 6 experimental conditions per tissue (bottom). **e-g**, For each experimental condition within each cell type, the percent of reads mapped to ribosomal genes (**e**), mitochondrial genes (**f**), and ERCC spike-ins (**g**) plotted against the mean number of genes expressed.

**Extended Data Fig. 2.**
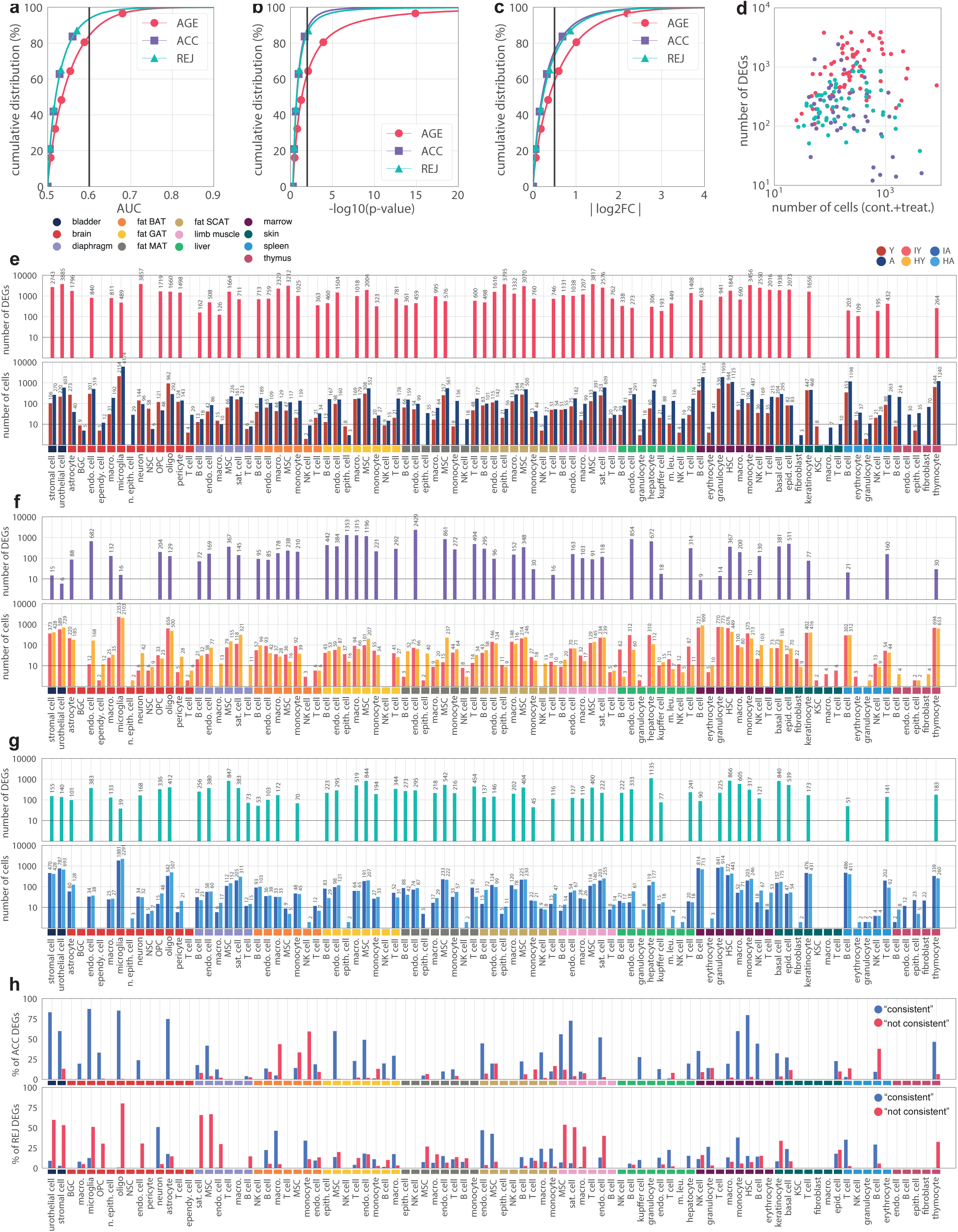
**a-c**, Cumulative distributions of the calculated AUC (**a**), -log_10_(p-value) (**b**) and log2 fold change values. Distributions are shown separately for ACC, REJ and AGE DGE. **d**, Number of DEGs plotted against the total number of cells within the control and treatment groups. Each dot represents a DGE comparison within a cell type. **e-g**, Number of DEGs (top) and sample sizes (bottom) of DGE comparisons for AGE (**e**), ACC (**f**) and REJ (**g**). **h**, Fraction of “consistent” and “not consistent” DEGs with AGE within the ACC (top) and REJ (bottom) comparisons.

**Extended Data Fig. 3.**
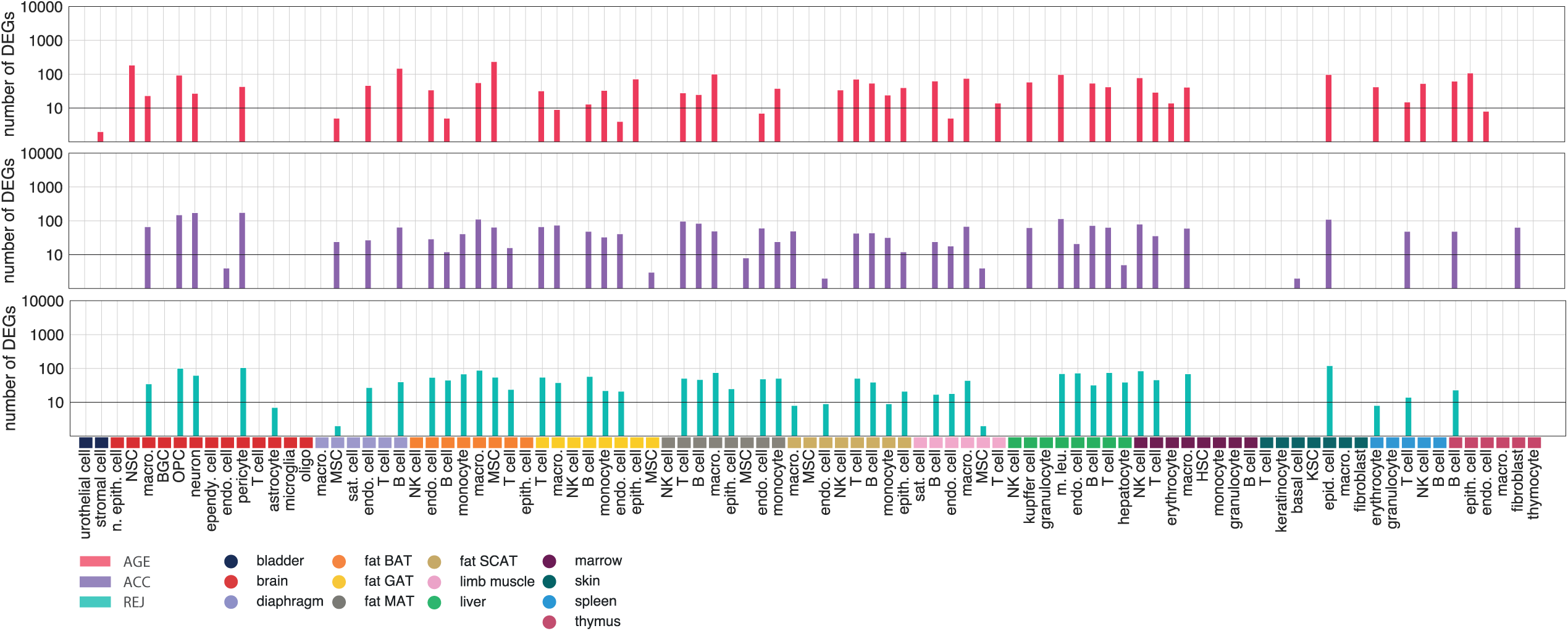
Number of DEGs shown for each AGE, ACC and REJ comparison after randomly permuting the condition labels of the cells within each cell type.

**Extended Data Fig. 4.**
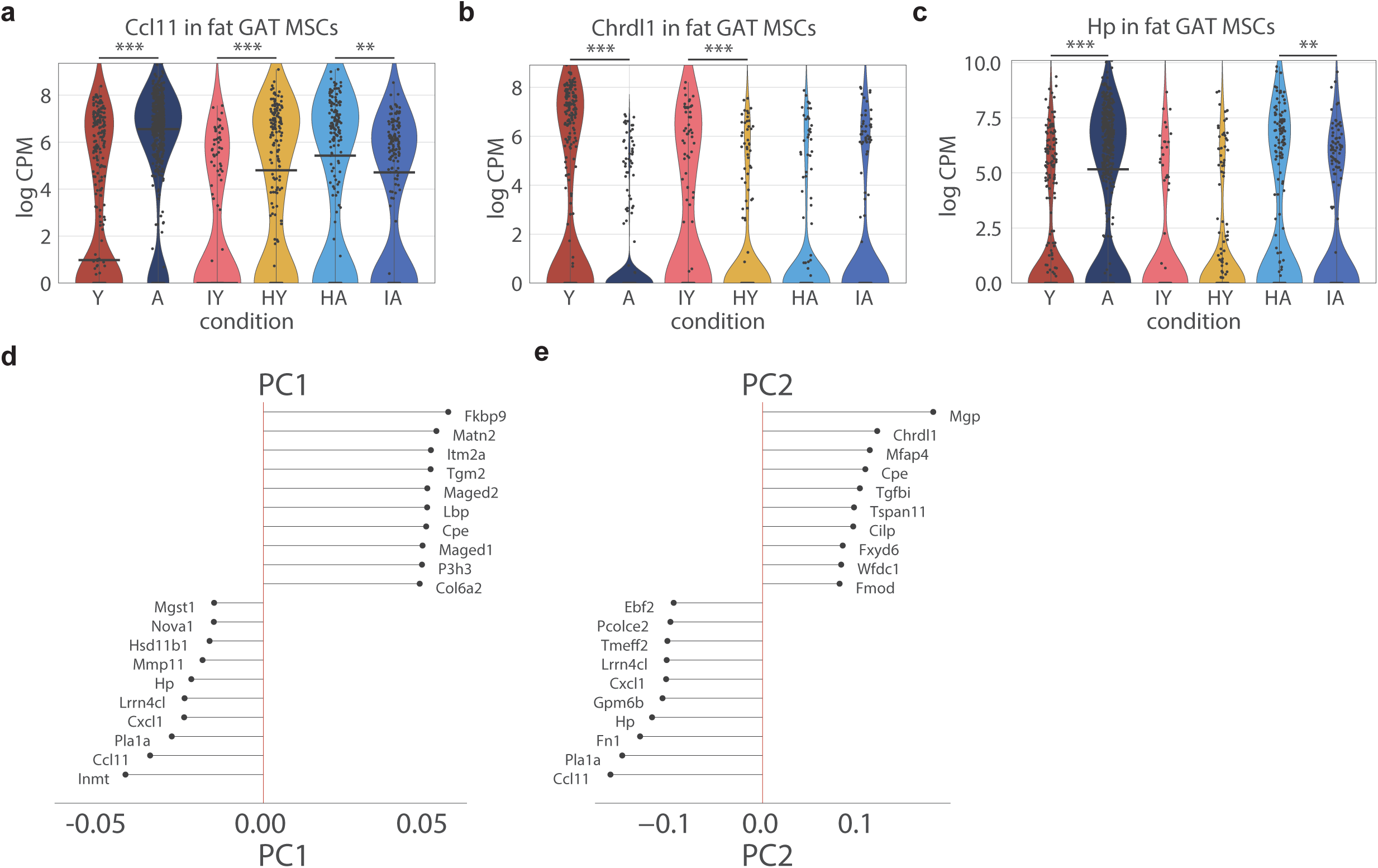
**a-c**, Violin plots showing the expression changes of Ccl11 (**a**), Chrld1 (**b**) and Hp (**c**) in fat GAT MSCs. **d-e**, Top genes associated with the first (**d**) and second (**e**) principal components within the fat GAT MSCs. PCA was conducted on DEGs after pre-selecting the strongest ageing genes (p<0.01, eff>0.6, |log_2_FC |>0.5).

**Extended Data Fig. 5.**
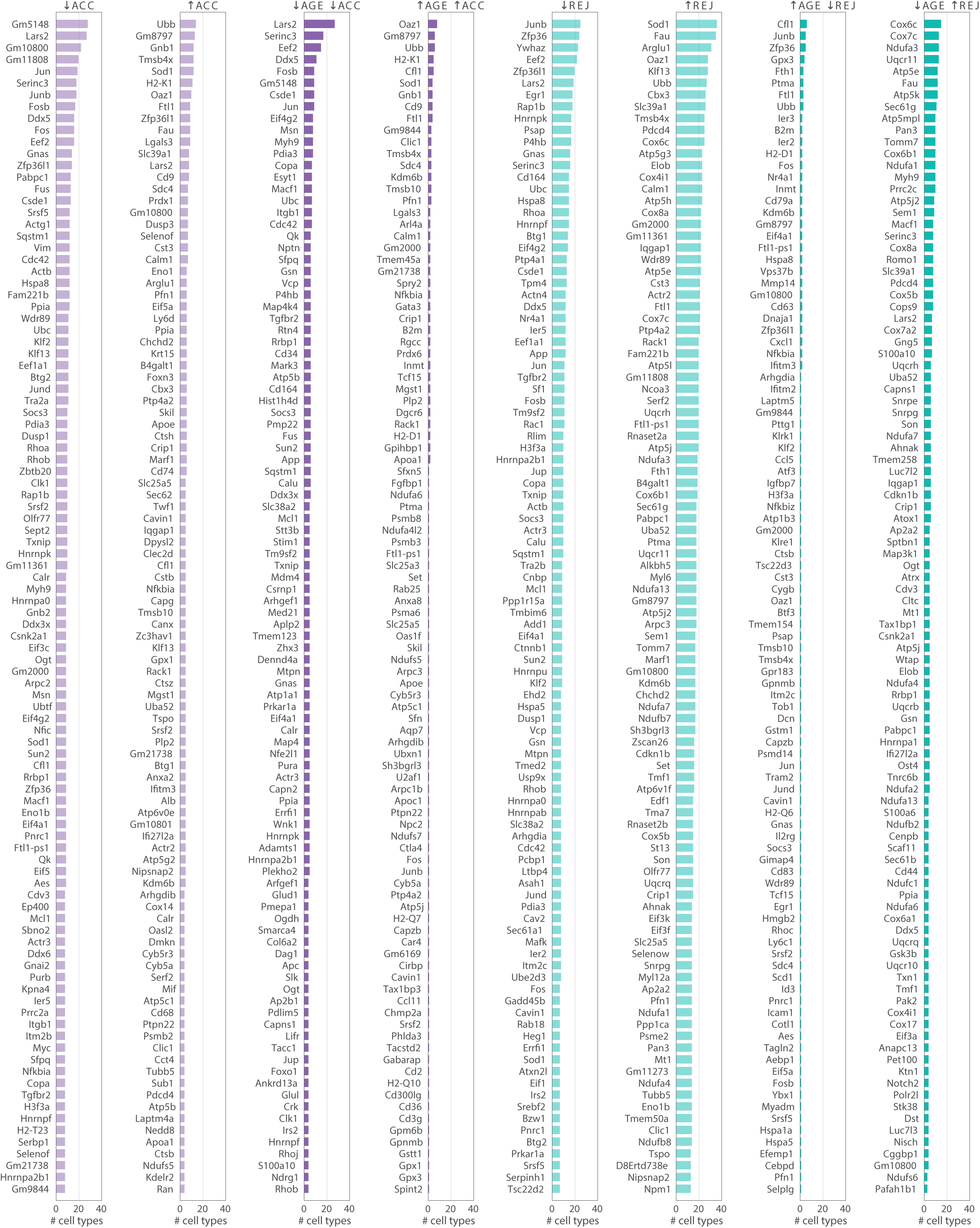
Top list of the 100 most frequent DEGs identified for ACC and REJ. Results are shown separately for up and downregulation. Columns with darker bars indicate top lists where only changes consistent with AGE are shown.

**Extended Data Fig. 6.**
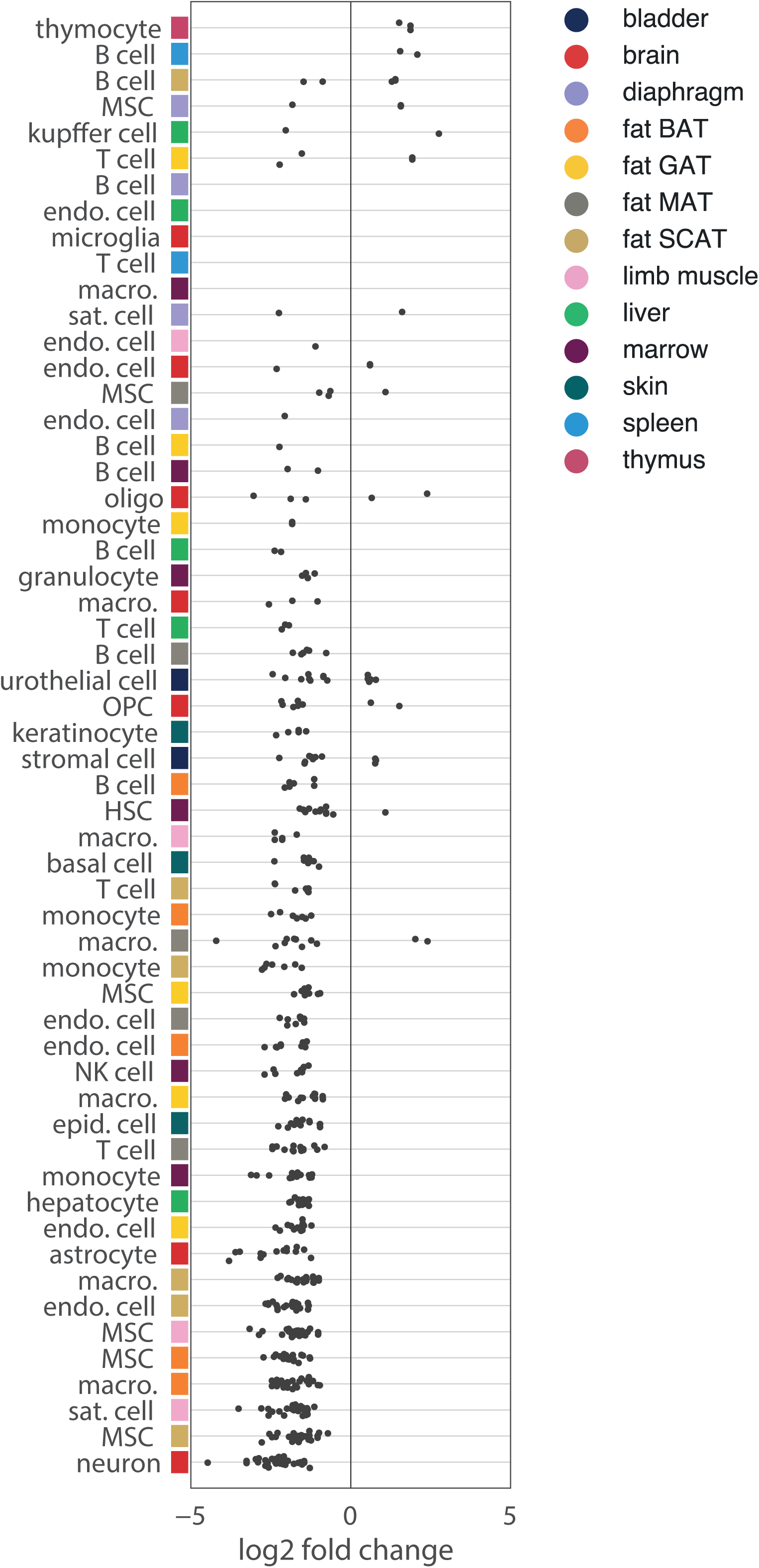
AGE log_2_FC fold changes of the 76 genes from the GO term “ATP synthesis coupled electron transport” (GO:0042773) within each cell type. Changes with (p<0.01, eff>0.6, |log_2_FC |>0.5) are indicated.

**Extended Data Table 1**. List of abbreviations used.

**Extended Data Table 2**. Number of cells per mouse and cell type.

**Extended Data Table 3**. List of cell type merging rules.

## The Tabula Muris Consortium

Nicole Almanzar^1^, Jane Antony^2^, Ankit S. Baghel^2^, Isaac Bakerman^2,3,4^, Ishita Bansal^2^, Ben A. Barres^5^, Philip A. Beachy^2,6,7,8^, Daniela Berdnik^9^, Biter Bilen^10^, Douglas Brownfield^6^, Corey Cain^11^, Charles K. F. Chan^12^, Michelle B. Chen^13^, Michael F. Clarke^2^, Stephanie D. Conley^14^, Spyros Darmanis^14^ ⍰, Aaron Demers^14^, Kubilay Demir^2,7^, Antoine de Morree^10^, Tessa Divita^14^, Haley du Bois^9^, Hamid Ebadi^14^, F. Hernán Espinoza^6^, Matt Fish^2,7,8^, Qiang Gan^10^, Benson M. George^2^, Astrid Gillich^6^, Rafael Gòmez-Sjöberg^14^, Foad Green^14^, Geraldine Genetiano^14^, Xueying Gu^8^, Gunsagar S. Gulati^2^, Oliver Hahn^10^, Michael Seamus Haney^10^, Yan Hang^8^, Lincoln Harris^14^, Mu He^15^, Shayan Hosseinzadeh^14^, Albin Huang^10^, Kerwyn Casey Huang^13,14,16^, Tal Iram^10^, Taichi Isobe^2^, Feather Ives^14^, Robert C. Jones^13^, Kevin S. Kao^2^, Jim Karkanias^14^, Guruswamy Karnam^17^, Andreas Keller^10,18^, Aaron M. Kershner^2^, Nathalie Khoury^10^, Seung K. Kim^8,19^, Bernhard M. Kiss^2,20^, William Kong^2^, Mark A. Krasnow^6,7^, Maya E. Kumar^21,22^, Christin S. Kuo^1^, Jonathan Lam^8^, Davis P. Lee^9^, Song E. Lee^10^, Benoit Lehallier^10^, Olivia Leventhal^9^, Guang Li^4,23^, Qingyun Li^5^, Ling Liu^10^, Annie Lo^14^, Wan-Jin Lu^2,6^, Maria F. Lugo-Fagundo^9^, Anoop Manjunath^2^, Andrew P. May^14^, Ashley Maynard^14^, Aaron McGeever^14^, Marina McKay^14^, M. Windy McNerney^24,25^, Bryan Merrill^16^, Ross J. Metzger^26,27^, Marco Mignardi^13^, Dullei Min^1^, Ahmad N. Nabhan^6^, Norma F. Neff^14^, Katharine M. Ng^6^, Patricia K. Nguyen^2,3,4^, Joseph Noh^2^, Roel Nusse^6,7,8^, Róbert Pálovics^10^, Rasika Patkar^17^, Weng Chuan Peng^8,38^, Lolita Penland^14^, Angela Oliveira Pisco^14^, Katherine Pollard^28^, Robert Puccinelli^14^, Zhen Qi^2^, Stephen R. Quake^13,14^⍰, Thomas A. Rando^9,10,29^, Eric J. Rulifson^8^, Nicholas Schaum^10^, Joe M. Segal^17^, Shaheen S. Sikandar^2^, Rahul Sinha^2,30,31,32^, Rene V. Sit^14^, Justin Sonnenburg^14,16^, Daniel Staehli^10^, Krzysztof Szade^2,33^, Michelle Tan^14^, Weilun Tan^14^, Cristina Tato^14^, Krissie Tellez^8^, Laughing Bear Torrez Dulgeroff^2^, Kyle J. Travaglini^6^, Carolina Tropini^16,39,40,41^, Margaret Tsui^17^, Lucas Waldburger^14^, Bruce M. Wang^17^, Linda J. van Weele^2^, Kenneth Weinberg^1^, Irving L. Weissman^2,30,31,32^, Michael N. Wosczyna^10^, Sean M. Wu^2,3,23^, Tony Wyss-Coray^9,10,29,34^⍰, Jinyi Xiang^1^, Soso Xue^13^, Kevin A. Yamauchi^14^, Andrew C. Yang^13^, Lakshmi P. Yerra^10^, Justin Youngyunpipatkul^14^, Brian Yu^14^, Fabio Zanini^13^, Macy E. Zardeneta^9^, Alexander Zee^14^, Chunyu Zhao^14^, Fan Zhang^26,27^, Hui Zhang^9^, Martin Jinye Zhang^35,36^, Lu Zhou^5^ & James Zou^14,35,37^

^1^Department of Pediatrics, Pulmonary Medicine, Stanford University School of Medicine, Stanford, CA, USA. ^2^Institute for Stem Cell Biology and Regenerative Medicine, Stanford University School of Medicine, Stanford, CA, USA. ^3^Stanford Cardiovascular Institute, Stanford University School of Medicine, Stanford, CA, USA. ^4^Division of Cardiovascular Medicine, Department of Medicine, Stanford University School of Medicine, Stanford, CA, USA. ^5^Department of Neurobiology, Stanford University School of Medicine, Stanford, CA, USA. ^6^Department of Biochemistry, Stanford University School of Medicine, Stanford, CA, USA. ^7^Howard Hughes Medical Institute, Chevy Chase, MD, USA. ^8^Department of Developmental Biology, Stanford University School of Medicine, Stanford, CA, USA. ^9^Veterans Administration Palo Alto Healthcare System, Palo Alto, CA, USA. ^10^Department of Neurology and Neurological Sciences, Stanford University School of Medicine, Stanford, CA, USA. ^11^Flow Cytometry Core, Veterans Administration Palo Alto Healthcare System, Palo Alto, CA, USA. ^12^Department of Surgery, Division of Plastic and Reconstructive Surgery, Stanford University, Stanford, CA, USA. ^13^Department of Bioengineering, Stanford University, Stanford, CA, USA. ^14^Chan Zuckerberg Biohub, San Francisco, CA, USA. ^15^Department of Physiology, University of California, San Francisco, CA, USA. ^16^Department of Microbiology & Immunology, Stanford University School of Medicine, Stanford, CA, USA. ^17^Department of Medicine and Liver Center, University of California San Francisco, San Francisco, CA, USA. ^18^Clinical Bioinformatics, Saarland University, Saarbrücken, Germany. ^19^Department of Medicine and Stanford Diabetes Research Center, Stanford University, Stanford, CA, USA. ^20^Department of Urology, Stanford University School of Medicine, Stanford, CA, USA. ^21^Sean N. Parker Center for Asthma and Allergy Research, Stanford University School of Medicine, Stanford, CA, USA. ^22^Department of Medicine, Division of Pulmonary and Critical Care, Stanford University School of Medicine, Stanford, CA, USA. ^23^Department of Developmental Biology, University of Pittsburgh School of Medicine, Pittsburgh, PA, USA. ^24^Mental Illness Research Education and Clinical Center, Veterans Administration Palo Alto Healthcare System, Palo Alto, CA, USA. ^25^Department of Psychiatry, Stanford University School of Medicine, Stanford, CA, USA. ^26^Vera Moulton Wall Center for Pulmonary and Vascular Disease, Stanford University School of Medicine, Stanford, CA, USA. ^27^Department of Pediatrics, Division of Cardiology, Stanford University School of Medicine, Stanford, CA, USA. ^28^Department of Epidemiology and Biostatistics, University of California, San Francisco, CA, USA. ^29^Paul F. Glenn Center for the Biology of Aging, Stanford University School of Medicine, Stanford, CA, USA. ^30^Department of Pathology, Stanford University School of Medicine, Stanford, CA, USA. ^31^Ludwig Center for Cancer Stem Cell Research and Medicine, Stanford University School of Medicine, Stanford, CA, USA. ^32^Stanford Cancer Institute, Stanford University School of Medicine, Stanford, CA, USA. ^33^Department of Medical Biotechnology, Faculty of Biochemistry, Biophysics and Biotechnology, Jagiellonian University, Krakow, Poland. ^34^Wu Tsai Neurosciences Institute, Stanford University School of Medicine, Stanford, CA, USA. ^35^Department of Electrical Engineering, Stanford University, Palo Alto, CA, USA. ^36^Department of Epidemiology, Harvard T.H. Chan School of Public Health, Boston, MA, USA. ^37^Department of Biomedical Data Science, Stanford University, Palo Alto, CA, USA. ^38^Princess Máxima Center for Pediatric Oncology, Utrecht, The Netherlands. ^39^School of Biomedical Engineering, University of British Columbia, Vancouver, British Columbia, Canada. ^40^Department of Microbiology and Immunology, University of British Columbia, Vancouver, British Columbia, Canada. ^41^Humans and the Microbiome Program, Canadian Institute for Advanced Research, Toronto, Ontario, Canada.

## References

1. Schaum, N. et al. Single-cell transcriptomics of 20 mouse organs creates a Tabula Muris. Nature 562, 367–372 (2018).

2. Schaum, N. et al. Ageing hallmarks exhibit organ-specific temporal signatures. Nature (2020). doi:10.1038/s41586-020-2499-y

3. Almanzar, N. et al. A single-cell transcriptomic atlas characterizes ageing tissues in the mouse. Nature 583, 590–595 (2020).

4. Castellano, J. M., Kirby, E. D. & Wyss-Coray, T. Blood-borne revitalization of the aged brain. JAMA Neurol. 72, 1191–1194 (2015).

5. Horvath, S. et al. Reversing age: dual species measurement of epigenetic age with a single clock. bioRxiv 2020.05.07.082917 (2020). doi:10.1101/2020.05.07.082917

6. Villeda, S. A. et al. The ageing systemic milieu negatively regulates neurogenesis and cognitive function. Nature 477, 90–96 (2011).

7. Katsimpardi, L. et al. Vascular and neurogenic rejuvenation of the aging mouse brain by young systemic factors. Science (80-.). 344, 630–634 (2014).

8. Smith, L. K. et al. β2-microglobulin is a systemic pro-aging factor that impairs cognitive function and neurogenesis. Nat. Med. 21, 932–937 (2015).

9. Khrimian, L. et al. Gpr158 mediates osteocalcin’s regulation of cognition. J. Exp. Med. 214, 2859–2873 (2017).

10. López-Otín, C., Blasco, M. A., Partridge, L., Serrano, M. & Kroemer, G. The hallmarks of aging. Cell 153, 1194 (2013).

11. Conboy, I. M. et al. Rejuvenation of aged progenitor cells by exposure to a young systemic environment. Nature 433, 760–764 (2005).

12. Chen, M. B. et al. Brain Endothelial Cells Are Exquisite Sensors of Age-Related Circulatory Cues. Cell Rep. 30, 4418–4432.e4 (2020).

13. Brigger, D. et al. Eosinophils regulate adipose tissue inflammation and sustain physical and immunological fitness in old age. Nat. Metab. 2, 688–702 (2020).

14. Ma, S. et al. Caloric Restriction Reprograms the Single-Cell Transcriptional Landscape of Rattus Norvegicus Aging. Cell 180, 984–1001.e22 (2020).

15. Das, M. M. et al. Young bone marrow transplantation preserves learning and memory in old mice. Commun. Biol. 2, 1–10 (2019).

16. Baht, G. S. et al. Exposure to a youthful circulaton rejuvenates bone repair through modulation of β-catenin. Nat. Commun. 6, 1–10 (2015).

17. Kovina, M. V., Zuev, V. A., Kagarlitskiy, G. O. & Khodarovich, Y. M. Effect on lifespan of high yield non-myeloablating transplantation of bone marrow from young to old mice. Front. Genet. 4, (2013).

18. Wang, C.-H. et al. Bone Marrow Rejuvenation Accelerates Re-Endothelialization and Attenuates Intimal Hyperplasia After Vascular Injury in Aging Mice. Circ. J. 77, 3045– 3053 (2013).

19. Smith, L. K. et al. The aged hematopoietic system promotes hippocampal-dependent cognitive decline. Aging Cell 19, (2020).

20. Pinti, M. et al. Aging of the immune system: Focus on inflammation and vaccination. European Journal of Immunology 46, 2286–2301 (2016).

## Methods References

21. Villeda, S. A. et al. Young blood reverses age-related impairments in cognitive function and synaptic plasticity in mice. Nat. Med. 20, 659–663 (2014).

22. Picelli, S. et al. Full-length RNA-seq from single cells using Smart-seq2. Nat. Protoc. 9, 171–181 (2014).

23. Darmanis, S. et al. A survey of human brain transcriptome diversity at the single cell level. Proc. Natl. Acad. Sci. U. S. A. 112, 7285–7290 (2015).

24. Picelli, S. et al. Tn5 transposase and tagmentation procedures for massively scaled sequencing projects. Genome Res. 24, 2033–2040 (2014).

25. Hennig, B. P. et al. Large-scale low-cost NGS library preparation using a robust Tn5 purification and tagmentation protocol. G3 Genes, Genomes, Genet. 8, 79–89 (2018).

26. Luecken, M. D. & Theis, F. J. Current best practices in single□cell RNA□seq analysis: a tutorial. Mol. Syst. Biol. 15, (2019).

27. Hagberg, A., Swart, P. & S. Chult, D. Exploring network structure, dynamics, and function using networkx. (2008). Available at: https://www.osti.gov/servlets/purl/960616.

28. Wolf, F. A., Angerer, P. & Theis, F. J. SCANPY: large-scale single-cell gene expression data analysis. Genome Biol. 19, (2018).

29. Pedregosa, F. et al. Scikit-learn: Machine Learning in Python. J. ofMachine Learn. Res. 12, 2825–2830 (2011).

30. McInnes, L., Healy, J. & Melville, J. UMAP: Uniform Manifold Approximation and Projection for Dimension Reduction. (2018).

31. Mann, H. B. & Whitney, D. R. On a Test of Whether one of Two Random Variables is Stochastically Larger than the Other. Ann. Math. Stat. 18, 50–60 (1947).

32. Gerstner, N. et al. GeneTrail 3: advanced high-throughput enrichment analysis. Nucleic Acids Res. 48, W515–W520 (2020).

33. Carbon, S. et al. The Gene Ontology Resource: 20 years and still GOing strong. Nucleic Acids Res. 47, D330–D338 (2019).

34. Kanehisa, M., Furumichi, M., Tanabe, M., Sato, Y. & Morishima, K. KEGG: New perspectives on genomes, pathways, diseases and drugs. Nucleic Acids Res. 45, D353– D361 (2017).

35. Benjamini, Y. & Hochberg, Y. Controlling the False Discovery Rate: A Practical and Powerful Approach to Multiple Testing. J. R. Stat. Soc. Ser. B 57, 289–300 (1995).

36. Gu, Z., Eils, R. & Schlesner, M. Complex heatmaps reveal patterns and correlations in multidimensional genomic data. Bioinformatics 32, 2847–2849 (2016).

37. Wickham, H. ggplot2. Wiley Interdiscip. Rev. Comput. Stat. 3, 180–185 (2011).

38. Csárdi, G. & Nepusz, T. *The igraph software package for complex network research*.

